# Contrasting determinants of mutation rates in germline and soma

**DOI:** 10.1101/117325

**Authors:** Chen Chen, Hongjian Qi, Yufeng Shen, Joseph Pickrell, Molly Przeworski

## Abstract

Recent studies of somatic and germline mutations have led to the identification of a number of factors that influence point mutation rates, including CpG methylation, expression levels, replication timing and GC content. Intriguingly, some of the effects appear to differ between soma and germline: in particular, whereas mutation rates have been reported to decrease with expression levels in tumors, no clear effect has been detected in the germline. Distinct approaches were taken to analyze the data, however, so it is hard to know whether these apparent differences are real. To enable a cleaner comparison, we considered a statistical model in which the mutation rate of a coding region is predicted by GC content, expression levels, replication timing, and two histone repressive marks. We applied this model to both a set of germline mutations identified in exomes and to exonic somatic mutations in four types of tumors. Germline and soma share most determinants of mutations; notably, we detected an effect of expression levels on germline mutations as well as on somatic ones. However, whereas in somatic tissues, increased expression levels are associated with greater strand asymmetry and *decreased* mutation rates, in ovaries and testes, increased expression leads to greater strand asymmetry but *increased* mutation rates. This contrast points to differences in damage or repair rates during transcription in soma and germline.

## Introduction

Germline mutations are the source of all heritable variation, including in disease susceptibility, and it is increasingly clear that somatic mutations also play important roles in human diseases, notably cancers (Muller 1927; Stratton, Campbell, and Futreal 2009). Understanding the rate and mechanisms by which mutations occur is therefore of interest to both evolutionary biologists and to human geneticists aiming to identify the underlying causes of genetic diseases (Shendure and Akey 2015; Gao et al. 2016). In particular, an accurate estimate of the local mutation rate is key to testing for an excess of disease mutations in specific genes among cases (Lawrence et al. 2013; Samocha et al. 2014). Characterization of the variation in mutation rate along the genome can also yield important insights into DNA damage and repair mechanisms (Stratton 2011; Ségurel, Wyman, and Przeworski 2014).

Until recently, our understanding of germline point mutations came mainly from analysis of diversity along the genome or divergence among species (Green et al. 2003; Webster et al. 2004; Polak and Arndt 2008; Hodgkinson and Eyre-Walker 2011; Park, Qian, and Zhang 2012). In the past several years, analyses have also been based on resequencing exomes or whole genomes from blood samples of human pedigrees and identifying variants present in the offspring but absent in the child (reviewed in Campbell and Eichler 2013 and Ségurel, Wyman, and Przeworski 2014; Shendure and Akey 2015; Francioli et al. 2015; Rahbari et al. 2016; Goldmann et al. 2016; Besenbacher et al. 2016). This approach is more direct than analyzing divergence data and presents the advantage of being almost unaffected by selection, but the analysis is technically challenging and, with current study designs, some mutations may be missed, notably those that occur in the early post-zygotic divisions (Rahbari et al. 2016; Moorjani, Gao, and Przeworski 2016; Harland et al. 2016).

Our knowledge of somatic point mutations, in turn, relies primarily on resequencing tumors. In these analyses, mutation calls are made by sequencing tumor and non-cancerous tissue pairs from the same individual and then excluding the variants shared between the two tissues (as the shared mutations are likely to be germline). Because, in this approach, a large population of cells is sequenced, the mutations identified tend to predate the tumorigenesis and thus are mostly somatic mutations that occurred in normal tissues (see, e.g., Martincorena et al. 2015; Alexandrov et al. 2015).

Studies of both germline and soma reveal that the point mutation rate varies across the genome, from the scale of a single base pair to much larger scales (Hodgkinson and Eyre-Walker 2011; Hodgkinson, Chen, and Eyre-Walker 2012; Ségurel, Wyman, and Przeworski 2014). At the single base pair level, the largest source of variation in germline mutation rate is the identity of the adjacent base pairs (Hwang and Green 2004; Hodgkinson and Eyre-Walker 2011). Notably, the mutation rate of CpG transitions (henceforth CpG Ti) is an order of magnitude higher than other mutation types (e.g., Kong et al. 2012). Most CpG dinucleotides are methylated in the human genome; when the methylated cytosine undergoes spontaneous deamination to generate thymine and is not corrected by the time of replication, the damage leads to a mutation. Among other types of sites, rates of mutation vary by 2 to 3 fold (Kong et al. 2012). In the soma, the mutation rate at CpG sites is also elevated, although the extent of the increase differs across tumor types (Pleasance, Stephens, et al. 2010; Pleasance, Cheetham, et al. 2010; Lee et al. 2010). More generally, tumors vary in their mutation spectrum: analyses of mutations and their two neighboring base pairs (i.e., considering 96 mutation types) point to enrichment of distinct mutational signatures for different types of cancers, a subset of which have been shown to reflect particular mutagens or differences in the efficiency of repair (Alexandrov et al. 2013).

Over a larger scale of megabases, germline mutation rates have been associated with a number of additional factors, including transcription level (in testis), replication timing (in lymphoblastoid cell lines), chromatin state (both in lymphoblastoid cells and in ovary), meiotic crossover rates and GC content (Hodgkinson and Eyre-Walker 2011; Michaelson et al. 2012; Park, Qian, and Zhang 2012; Francioli et al. 2015; Goldmann et al. 2016; Besenbacher et al. 2016). Somatic mutation rates have also been associated with replication timing (in Hela cell lines) and with average transcription levels across 91 cell lines in Cancer Cell Line Encyclopedia (Lawrence et al. 2013).

In many cases, little is known about the mechanistic basis for the association of a given factor with mutation rates. However, the association of somatic mutation rates with transcription levels appears to be a byproduct of transcription-coupled repair (TCR), a sub-pathway of nucleotide excision repair (NER) (Hanawalt and Spivak 2008; Nouspikel 2009). NER is a versatile repair pathway that senses lesion-causing distortions to DNA structure and excises the lesion for repair. Another subpathway of NER, global genome repair (GGR), can repair lesions on both transcribed strand (henceforth TS) and non-transcribed strand (henceforth NTS), including transcribed regions as well as transcriptionally-silent ones. In contrast, TCR operates only within transcribed regions, triggered by lesions on the TS, which it repairs off the NTS. This mechanism gives rise to a mutational strand asymmetry as well as a compositional asymmetry between strands. For example, TCR leads to more A to G mutations (A>G henceforth) on the NTS than TS; acting over long periods of time, this phenomenon generates an excess of G over A (and T over C) on the NTS (Green et al. 2003; McVicker and Green 2010). Such mutational strand asymmetry has been found in both germline and soma (Green et al. 2003; Polak and Arndt 2008; Rubin and Green 2009; Lawrence et al. 2013; Martincorena et al. 2015; Francioli et al. 2015).

While many of the same determinants appear to play important roles in both germline and soma, there are hints of differences as well. For instance, studies of pre-neoplastic somatic mutations indicate that, over a 100 kb scale, the mutation rates in somatic tissues decrease with expression levels and increase with replication timing (Lawrence et al. 2013). Similarly, two studies that focused on somatic mutations in non-cancerous somatic tissues, normal eyelid tissue and neurons, found mutations to be enriched in regions of low expression and repressed chromatin (Martincorena et al. 2015; Lodato et al. 2015). A similar effect of replication timing was identified in studies of germline mutation (Stamatoyannopoulos et al. 2009; Francioli et al. 2015; Besenbacher et al. 2016; Carlson et al. 2017). However, the effect of expression levels on germline mutation rates remains unclear: one study reported increased divergence between humans and macaques with greater germline expression (Park, Qian, and Zhang 2012), but others found no discernable effect of expression levels on mutation rates (Green et al. 2003; Webster et al. 2004; Polak and Arndt 2008; Hodgkinson and Eyre-Walker 2011; Francioli et al. 2015). This difference between germline and soma is particularly puzzling in light of the observation that the strand asymmetry of mutation rates between TS and NTS is seen in the germline as well as the soma (Pleasance, Cheetham, et al. 2010; Pleasance, Stephens, et al. 2010; McVicker and Green 2010; Lawrence et al. 2013). Together, these observations suggest that the determinants of mutation rates may differ between germline and soma, raising the more general possibility that the damage rate or the repair efficacy differs among cell types (Lynch 2010).

A limitation, however, is that studies have used different statistical approaches, rendering the comparison hard to interpret. As an illustration, whereas some studies binned the genome into windows of 100 kb (e.g., Lawrence et al. 2013) or 1Mb regions (e.g., Polak et al. 2015), other studies have compared the mean mutation rate in transcribed regions and non-transcribed regions or in genes grouped by expression levels (Hodgkinson and Eyre-Walker 2011; Francioli et al. 2015; Lodato et al. 2015). Studies of somatic mutation also vary in whether they group different tissues or distinguish among them (e.g., Pleasance, Stephens, et al. 2010; Lawrence et al. 2013). An additional limitation of earlier studies of germline mutation is that, by necessity, they relied on human-chimpanzee divergence as a proxy for de novo mutation rates (Green et al. 2003; Webster et al. 2004; Hodgkinson and Eyre-Walker 2011), even though divergence reflects not only the mutation process but also effects of natural selection in the human-chimpanzee ancestor and biased gene conversion (McVicker et al. 2009; Duret and Galtier 2009).

To our knowledge, only one study has used a uniform approach to study germline and soma. Their findings point to possible differences in their determinants: for instance, the histone mark H3K9me3 accounts for more than 40% of mutation rate variation at 100 kb in tumors, when a much weaker association is seen in the germline (Schuster-Böckler and Lehner 2012; Goldmann et al. 2016). This analysis relied on pairwise correlations, however, and therefore the results may be confounded by other factors that are correlated to the histone marks and differ between tissues. Moreover, to our knowledge, there has been no parallel treatment of strand asymmetry in germline and soma.

To overcome these limitations, we built a multivariable regression model, in which the mutation rates of CpG Ti and other types of mutations in a coding region are predicted by GC content, expression levels, replication timing and two histone repressive marks. To this end, we used the expression levels, replication timing and histone marker levels of matched cell types. We applied the model to a large set of germline point mutations identified in exomes from recently published studies on developmental disorders and to somatic point mutations in exomes found in four types of tumors and reported by the Cancer Genome Atlas (see Materials and Methods). In addition, we considered the mutational strand asymmetry in the two sets of data.

## Materials and Methods

### Datasets

To study germline mutations, we relied on de novo mutation calls made from 8681 trios surveyed by exome sequencing. We combined results from two main sources: studies of neurodevelopmental disorders (NDD), which considered 5542 cases and 1911 controls (unaffecteds), and studies of congenital heart defect (CHD), conducted by the Pediatric Cardiac Genomics Consortium, which included 1228 trios. The NDD cases include 3953 cases of Autism Spectrum Disorder (ASD), 1133 cases of deciphering developmental disorders (DDD), 264 cases of epileptic encephalopathies (EE), and 192 cases of intellectual disability (ID). All these studies applied similar capture and sequencing methods, and most samples were at >20X coverage (see Table 1). We tested for an effect of the study, which could potentially arise from differences in design or analysis pipeline, by adding a categorical variable (by an analogous approach to the one described below to test for differences among tissues). We found a marginally significant interaction between the study and the expression level in testis (our proxy for expression levels in the germline), driven by one study (CHD cases; Homsy et al. 2015), as well as for interactions between the studies and the effects of H3K9me3 and GC content, driven by two small studies (EE and ID) (see Figure S1). Given these very minor differences and in order to increase our power, we combined all the germline mutation datasets in what follows (see Supplementary Materials Table S1 for list of mutations).

**Table 1.** Summary of germline datasets

| Datasets | Trios | References | Capture | Sequencing |
| --- | --- | --- | --- | --- |
| Autism Spectrum Disorder (ASD) | 3953 | De Rubeis et al. 2014; Iossifov et al. 2014 | Exome | Illumina and SOLiD |
| Simons Simplex Collection, unaffected | 1911 | Iossifov et al. 2014 | Exome | Illumina |
| Congenital heart disease (CHD) | 1213 | Homsy et al. 2015 | Exome | Illumina |
| Deciphering Developmental Disorders Study (DDD) | 1133 | The Deciphering Developmental Disorders Study 2015 | Exome | Illumina |
| Epileptic Encephalopathies (EE) | 264 | Epi4K Consortium and Epilepsy Phenome/Genome Project 2013 | Exome | Illumina |
| Intellectual Disability (ID) | 192 | de Ligt et al. 2012; Rauch et al. 2012; Hamdan et al. 2014 | Exome | Illumina |

To examine determinants of mutation rates in somatic tissues, we downloaded somatic mutation calls identified in four types of cancer from the Cancer Genome Atlas (TCGA) portal (in July 2015): breast invasive carcinoma (BRCA), cervical squamous cell carcinoma and endocervical adenocarcinoma (CESC), brain lower grade glioma (LGG), and liver hepatocellular carcinoma (LIHC). The numbers of samples are listed below (Table 2). In all cases, both non-cancerous and tumor tissues of patients were sampled and the exomes were sequenced using an Illumina platform. In the studies, mutation calls shared by the normal and tumor samples were removed (on the presumption that they are germline). What remains are somatic mutations found at high enough frequency to be seen in a large population of cells, which are therefore likely to predate the tumorigenesis, i.e., mutations that occurred in the pre-neoplastic tissues (Martincorena et al. 2015).

**Table 2.** Sizes of TCGA datasets

| Datasets | Sample sizes |
| --- | --- |
| Breast Invasive carcinoma (BRCA) | 904 |
| Cervical squamous cell carcinoma and endocervical adenocarcinoma (CESC) | 181 |
| Low grade glioma (LGG) | 502 |
| Liver hepatocellular carcinoma (LIHC) | 171 |

For each type of cancer with more than one mutation annotation file available in the TCGA data portal, we selected the file that included the largest number of patient samples. We removed the ~7.6% of samples that had an unusually large number of mutations per sample (p<0.05 by Tukey’s test), because they are likely to reflect loss of some aspect of the DNA mismatch repair and hence their mutational mechanisms likely differ (Supek and Lehner 2015).

### Possible determinants of mutation rates

We considered the main factors previously reported to be significantly correlated with mutation rates, namely expression levels, replication timing, GC content and histone modification levels. To quantify expression levels, we relied on gene expression data (measured as RPKM) from the Genotype-Tissue Expression (GTEx) for breast, uterus, brain cortex and liver tissues. We used gene expression levels of testis and ovary as our proxy for germline expression.

The effect of the replication timing on somatic mutation rates was argued to be cell-type specific (Supek and Lehner 2015). We therefore relied on Repli-Seq measurements (provided per base pair) in ENCODE cell lines that match the four types of cancer, namely MCF-7 (breast cancer), Hela-S3 (cervical cancer), SK-N-SH (neuroblastoma), and HepG2 (liver hepatocellular carcinoma) cell lines.

These measurements were obtained from the UCSC Genome Browser. In all cases, the replication timing reported is a smooth measure of the relative enrichment of early vs. late S-phase nascent strands, with high values indicating early replication. For each gene, we computed the average replication timing by taking the mean value of the data points that overlap with gene start-to-end coordinates in UCSC Refseq gene database. For genes with multiple transcripts, we took the union of all exons in all transcripts. For germline mutations, there are no data for the appropriate cell types, so we used replicating timing estimates for lymphoblastoid cell lines (LCL) (provided in 10 kb windows) (Koren et al. 2012). We also tried using replication timing data from three somatic tissues instead; the replication timing data are highly correlated among the tissues and therefore the effects of mutation were estimated to be very similar (see Figure S2).

We also considered the effects of chromatin marks that had been shown to correlate individually with somatic and germline mutation rates (Schuster-Böckler and Lehner 2012; Carlson et al. 2017): specifically, histone modification H3K9me3 and H3K27me3, two repressive marks associated with constitutively and facultatively repressed genes, respectively. Levels of these marks were downloaded from roadmap epigenomics data browser (Dec 2015, hg19) and converted to gene-based histone modification levels by averaging across the gene. We used the histone modification levels of adult ovary, breast myoepithelial cells, brain hippocampus and adult liver as proxies for germline, breast, brain and liver, respectively. In the following regression analysis, we considered only three of four somatic tissues, as we could not obtain histone modification data for CESC. Finally, we computed exonic GC content as the fraction of G or C residues in the union of exons in all isoforms of a given gene.

Germline mutation studies relied on the UCSC Refseq gene annotation, whereas TCGA uses GENECODE annotation, which contains more transcripts (Larsson et al. 2005; Zhao and Zhang 2015). To make the comparison cleaner, we focused on exonic regions considered in both types of studies by using gene and exon coordinates of Refseq database in build hg19 from UCSC genome browser.

### Statistical model

Our main goal was to investigate possible relationships between mutation rates and gene expression levels, while controlling for replication timing, GC content and some histone modification levels. Because our mutation counts are over-dispersed, with greater variance than mean, we used a negative binomial regression model (instead of, e.g., a Poisson regression model). Specifically, for every protein-coding gene, we counted the number of CpG Ti or other types of mutations in the coding exons of a gene and treated it as an outcome of a sequence of independent Bernoulli trials with probability λ_i_ where λ_i_ is the probability of a mutation occurring in gene i.

Transitions at CpG sites are thought to primarily occur due to spontaneous deamination at methylated cytosines, a distinct mutational source, and thus their determinants may be distinct from other mutation types (reviewed in Ségurel, Wyman, and Przeworski 2014). However, within CpG islands, most CpGs are hypomethylated (Takai and Jones 2002). To focus on a more homogeneous set of methylated CpGs, we therefore excluded CpG islands from the analyses of CpG Ti. CpG island annotations were downloaded from UCSC browser (track: CpG Islands).

We considered gene expression levels measured in RPKM (X_1_), replication timing (X_2_), mean histone modification levels (H3K9me3 as X_3_, H3K27me3 as X_4_) and GC content (X_5_) as predictors. We also included L, the total number of CpG sites (when considering CpG Ti) or all nucleotides (when considering all other types of mutations) in the exons of the given gene, as an exposure variable, to account for the variation in gene length. The logarithm of λ_i_ is then modeled as a linear combination of these features scores:

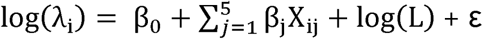

We used R function glm.nb to estimate the coefficients, where β_0_ is an intercept term, β_j_ is the effect size of feature j, and X_ij_ is the score for feature j in gene i. In order to make the effect sizes of different features comparable within a model, we normalized all the predictor variables to have a mean of 0 and a standard deviation of 1. The gene expression levels measured in RPKM originally range from 0 to a few hundred thousand. As is standard (e.g., Green et al. 2003; Francioli et al. 2015), we added half of the smallest non-zero value in the corresponding expression data sets and then log-transformed the expression level before normalization.

We note that in this model, we are considering possible effects one at a time. Including interaction terms affects the estimates and significance levels but changes none of the qualitative results, with the exception of results for H3K27me3, which become less significant (see Figure S3).

To examine whether the predictors have significantly different effects across tissues, we combined the models into one by including a categorical variable C for the tissue type (see Figure 2). In this approach:

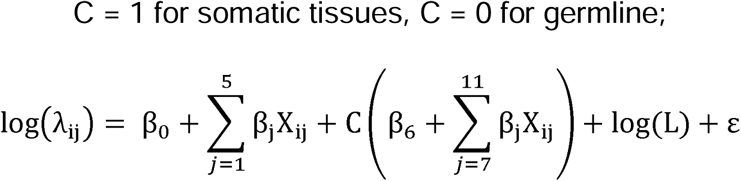

X_1_, X_2_, X_3_, X_4_ and X_5_ are the same genomic or epigenomic features as in the separate model, **β**_1_, **β**_2_, **β**_3_, **β**_4_, **β**_5_ are the effect sizes of features X_1_ to X_5_ for testis, and **β**_7_, **β**_8_, **β**_9_, **β**_10_, **β**_11_ are the differences of effect size in the somatic tissue of features X_1_ to X_5_ compared to those in testis. We used the R function glm.nb to estimate the coefficients.

Similarly, in order to ask whether effects differ between CpG Ti and other type of mutations in the same tissue, we included a binary variable C for the two mutation types (see Figure S4).

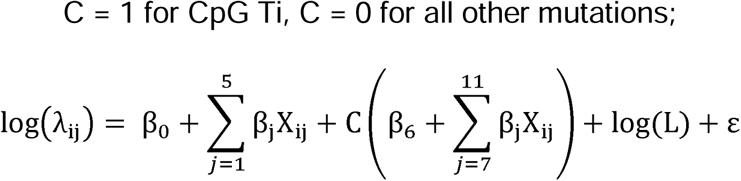

All variables are set up the same way as in the combined model described previously, except for that **β**_7_, **β**_8_, **β**_9_, **β**_10_, **β**_11_ are now the differences of the effect sizes for CpG Ti compared to those for all other mutation types.

### Mutation spectrum and strand asymmetry analysis

We annotated the direction of transcription using the UCSC CCDS track and filtered out genes that are transcribed off both strands (1.7% of genes in Refseq), which left around 19,000 genes to consider. This annotation allowed us to classify mutations into six types of mutation (A>C, A>G, A>T, G>A, G>C, G>T) on either TS or NTS. There are thus 12 possible changes (each of the six on both strands). We then calculated the mutation rate of any given type on NTS and TS separately, by considering the number of corresponding mutations in the combined data sets, divided by the total number of nucleotides that could give rise to such a mutation in the exons. To obtain the confidence intervals on the mutation rates (reported in Figure 3, 4 and Figure S5) as well as for the mutation asymmetry ratio (Figure 4 and Figure S5), we used bootstrap. Specifically, we created 100 samples, of the same size as the original sample, by drawing randomly from the original sample with replacement, and estimated the 95% CI from those 100 samples.

We tested for strand asymmetry by a Chi-squared test. Because A>G strand asymmetry shows the greatest asymmetry (Green et al. 2003) and is the only mutation type that we found in all tissues (Figure 3), we focused primarily on this type, though we also considered A>T mutational patterns (see Figure S5). To test if the extent of strand asymmetry changes with transcription levels, we grouped genes into expression level quantiles and calculated A>G strand asymmetry. Our measure of strand asymmetry is the ratio of the mutation rate on NTS to that on TS.

### Data availability

Germline mutations are provided in Table S1. TCGA somatic mutations can be downloaded from GDC data portal (https://gdc-portal.nci.nih.gov/search/s?facetTab=cases). The gene RPKM data are available at GTEx website (http://www.gtexportal.org/home/datasets). The replication timing data of LCL and other tissues are available from (Koren et al. 2012) and ENCODE website (https://www.encodeproject.org/search/?type=Experiment&assay_title=Repli-seq) respectively. The histone modification data can be freely accessed at epigenome roadmap website (http://www.roadmapepigenomics.org/data/tables/all).

## Results

We began by applying our multivariable regression model (see Materials and Methods) to compare the determinants of mutation rates per gene between the two germline tissues and among the three somatic tissues (Figure 1). Results for germline mutations are very similar using testis or ovary expression profiles. In both, there is no discernable effect of replication timing, other than a marginally significant negative effect for mutations other than CpG Ti. However, in contrast to a previous study using de novo mutations (Francioli et al. 2013) and most previous studies of divergence, we found a significant increase of germline mutation rates with expression levels for both CpG Ti and other mutation types (Figure 1; see also Figure S2 for similar results with replication timing for different tissues). The difference with a previous analysis of de novo mutations may be due to the scale of a gene considered here (rather than 100 kb windows).

**Figure 1.**
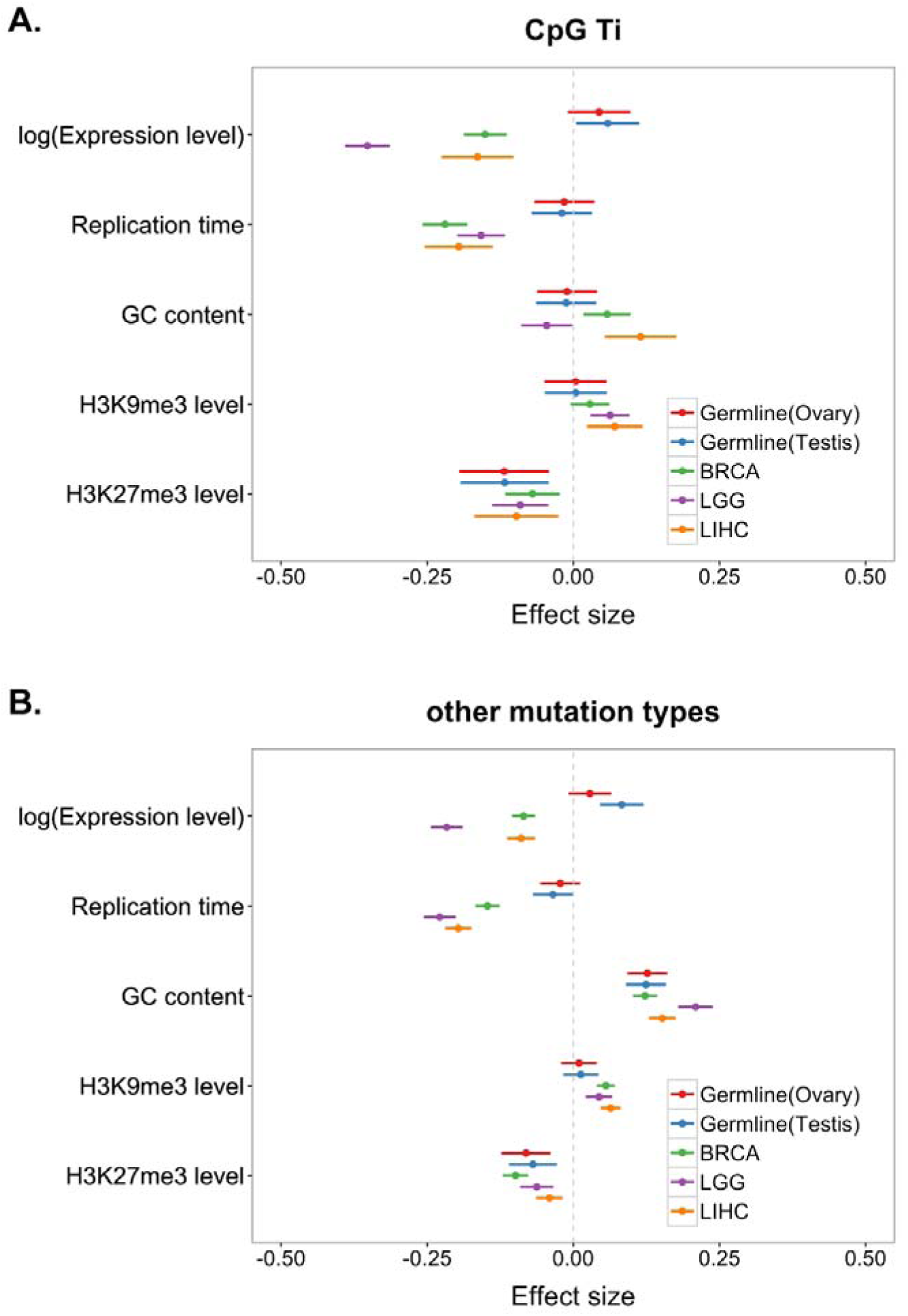
Coefficients of multivariable binomial regression model fit to germline and somatic mutation data. In panel A, are results for CpG Ti and in panel B, for other mutation types. Red, blue and green, purple and orange bars represent the 95% CI for the estimate of the regression coefficient in germline data set using ovary expression and testis expression, BRCA (breast invasive carcinoma), LGG (brain lower grade glioma) and LIHC (liver hepatocellular carcinoma) data sets respectively. For all replication timing data, high value means early.

The effect of expression levels is most clearly seen using testis expression (P = 0.03 for CpG Ti; P = 1.4x10^−5^ for other mutation types) than using ovary expression, possibly due to the fact that over three quarters of germline mutations are of male origin (Kong et al. 2012; Rahbari et al. 2016; Goldmann et al. 2016). Alternatively, the ovary expression profile may be a poorer proxy for female germ cells than the testis expression profile is for male germ cells. In any case, henceforth, we use testis expression profile for analysis of the germline mutation rates.

We note that our analysis of germline mutation relies on calls made in exome studies of blood samples from six sets, including five cases and unaffected controls (see Table 1). A previous study reported that in one set of cases, individuals with congenital heart disease (CHD), there is an increased number of putatively damaging mutations in the genes most highly expressed in the developing heart and brain (Homsy et al. 2015). Since the mutations are thought to be germline mutations (rather than somatic mutations), this association cannot be causal, instead reflecting an enrichment of damaging mutations in important heart developmental genes in CHD patients. To evaluate whether our findings of increased mutation rates with germline expression levels could be driven by a similar ascertainment bias, we excluded the CHD set and obtained the same results (see Figure S6). We also reran the analysis, comparing the effects in the five cases compared to the controls; none of the qualitative results differed, though as expected from the smaller size of the control sets, the estimated effect sizes were more uncertain (see Figure S7). Thus, our results suggest that the increase in mutation rates with expression levels in testes is not a result of focusing primarily on cases.

Germline mutation rates are also associated with H3K27me3 levels. We also found that, other than for CpG Ti, mutation rates in a gene increase with its GC content. This observation is consistent with previous findings of a high rate of GC to AT mutations relative to other types (e.g., Kong et al. 2012). Moreover, it is thought that mis-incorporated bases during DNA replication in an AT rich regions are more easily accessible and thus more easily repaired than GC rich regions (Petruska and Goodman 1985; Bloom et al. 1994). In contrast, we found a marginally negative effect of GC content on germline rates of CpG Ti. A possible explanation for this observation is that spontaneous deamination, the likely source of most CpG Ti, occurs more readily when DNA is single stranded, which is more likely in AT-rich than GC-rich regions (Fryxell and Moon 2005; Elango et al. 2008).

The effects of determinants on mutation rates are also concordant across somatic tissues. Notably, mutation rates decrease with expression levels in all three tissues, though the magnitudes of the effects differ. This finding is consistent with previous studies and thought to be a result of TCR (Lawrence et al. 2013). Intriguingly, in a model comparing the effects on CpG Ti and other mutation types directly, in all three somatic tissues, the effect of expression levels on mutation rates is most pronounced for CpG Ti (see Figure S4). This finding suggests that damage or repair of CpG Ti is tightly coupled to transcription.

In all three somatic tissues, there is also a decrease in mutation rate with replication timing and H3K27me3 levels, as well as an increase with H3K9me3 levels (Schuster-Böckler and Lehner 2012; Behjati et al. 2014; Blokzijl et al. 2016). The effect of replicating timing on mutation rate has been attributed to the depletion of free nucleotides within later replicating regions, leading to the accumulation of single-stranded DNA and thus rendering the DNA more susceptible to endogenous DNA damage (Stamatoyannopoulos et al. 2009). An alternative hypothesis is that DNA mismatch repair (MMR), which is coupled with replication, is more effective in the early replicating regions of the genome; this possibility is supported by the finding that this association is not detected in the tissue of MMR-deficient patients (Supek and Lehner 2015). While on face value, it may seem surprising that replication timing is a significant determinant for the LGG samples, given that neurons are post-mitotic, glial cells still retain their ability to divide and a substantial fraction of mutations detected in neuronal samples may have occurred at earlier stages in development.

The only difference in the determinants of mutation rates across somatic tissues appears to be the effect of GC content on CpG Ti rates: mutation rates decrease with GC content in brain tissues and increase with GC content in liver and breast tissues. This finding raises the possibility that damage or repair rates of CpG sites differ in brain tissues (Lodato et al. 2015).

Figure 1 also hints at a difference between testes (also ovaries) and somatic tissues in the directional effect of expression levels on mutation rates, with a marginally significant positive effect for germline mutations (P = 0.03 for CpG Ti, P = 1.4x10^−5^ for other mutation types) and a significantly negative effect for somatic tissues (e.g., BRCA: P = 8x10^−16^ for CpG Ti; P<2x10^−16^ for other mutation types). When we tested for this difference explicitly, by adding a binary variable for soma and germline (see Materials and Methods), we found that expression levels and replication timing differ in their effects, for both CpG Ti and other mutation types (Figure 2).

**Figure 2.**
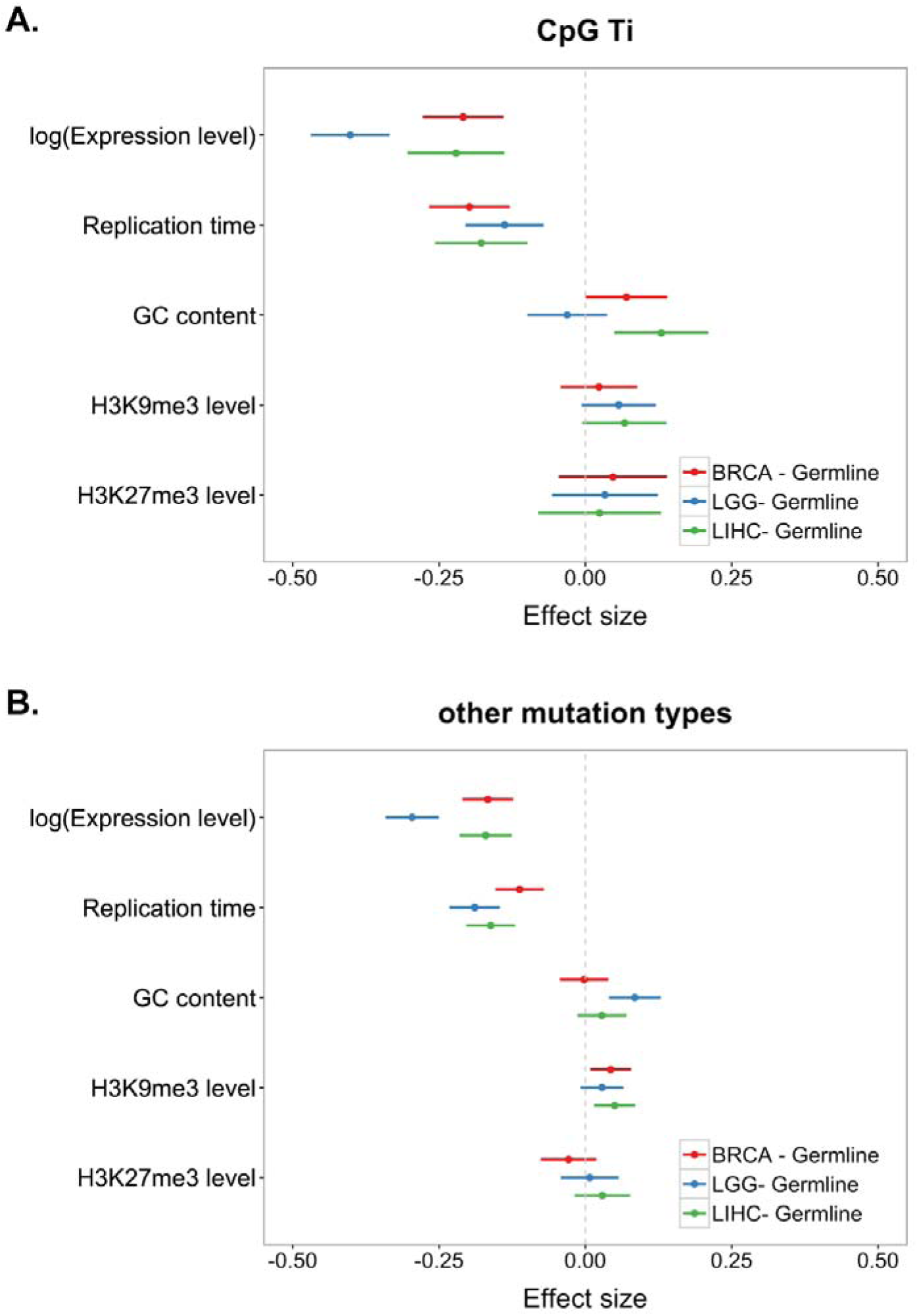
Coefficients of combined model comparing each somatic data set to germline data set using testis expression. In panel A, results for CpG Ti and in panel B, for other mutation types. Red, blue and green bars represent the 95% CI of the deviation of the estimated coefficient from the germline estimate; they are shown for BRCA (breast invasive carcinoma), LGG (brain lower grade glioma) and LIHC (liver hepatocellular carcinoma) data sets respectively. For all replication timing data, high value means early.

Specifically, replication timing has a positive effect on both tissue types but its effect is stronger in soma (Figure 2). The simplest explanation is that a larger fraction of mutations in the soma are introduced by errors related to replication, as opposed to other non-replicative sources. Another (not mutually-exclusive) possibility is that the effect of early replication versus late replication differs to a greater extent in the soma than in the germline. For example, if MMR is much more efficient in early replicating regions (Supek and Lehner 2015) and more efficient in soma than germline.

To examine this possibility further, we considered a signature of TCR—strand asymmetry—in the different tissues, finding it among germline mutations as well as in all four somatic tissues (Figure 3). Consistent with previous studies (Green et al. 2003; Francioli et al. 2015), one type in particular, A > G, stands out. While the asymmetry is significant in all five data sets, with more mutation on the NTS than the TS, the degree of asymmetry is significantly different among the five data sets (χ^2^ test, P = 3x10^−8^), with the strongest seen in germline. Intriguingly, other mutation types, notably G>C mutations, show even more pronounced differences among tissues, with significant excess on the transcribed strand in the germline and LGG samples but a significant paucity on the NTS in BRCA and CESC. These findings indicate a potential difference in either strand-biased damage or in TCR (or both) among somatic tissues. In summary, the total mutation rate appears to behave quite differently as a function of expression levels in the germline and the soma (Figure 1 and 2), despite the fact that we observed clear evidence for TCR in both types of tissues (Figure 3).

**Figure 3.**
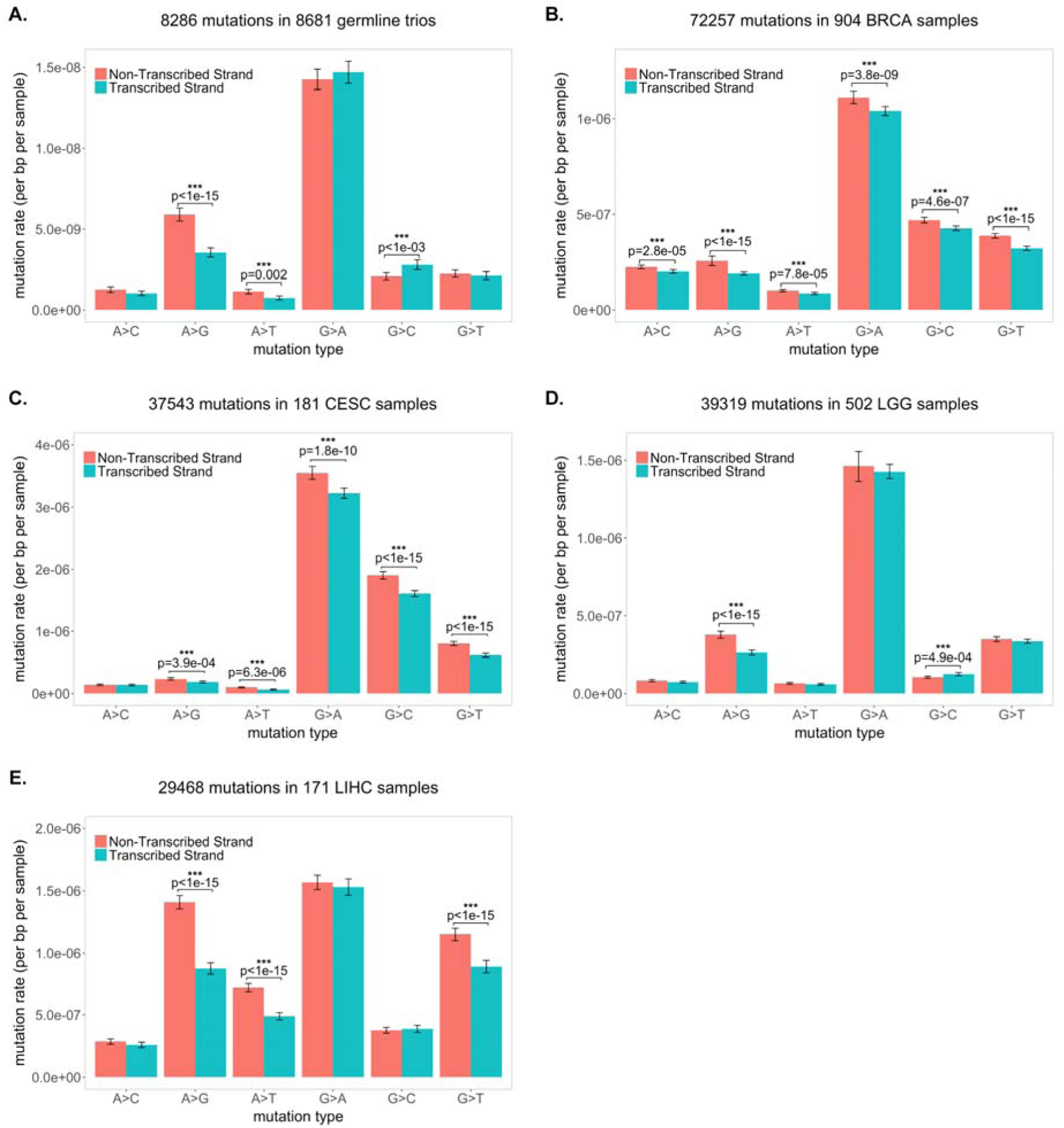
Strand asymmetry for six mutation types. In panel A are results for the germline; in panel B, for BRCA (breast invasive carcinoma); in panel C, for CESC (cervical squamous cell carcinoma and endocervical adenocarcinoma); in panel D, for LGG (brain lower grade glioma); and in panel E, for LIHC (liver hepatocellular carcinoma). The error bars of the mutation rate denote 95% confidence intervals estimated by bootstrapping (see Materials and Methods).

To examine this difference in more detail, we focused on A>G mutations and considered how the mutation rate and degree of asymmetry covary with expression (Figure 4). A striking contrast emerges: in the germline, as expression levels increase, mutation rates and asymmetry increase, whereas in the soma, asymmetry increases while mutation rates decrease. The same pattern is seen when A>T mutation rate and asymmetry are considered (see Figure S5). This difference in behavior with expression levels strongly suggests that the balance between damage and repair rates during transcription differs between germline and soma.

**Figure 4.**
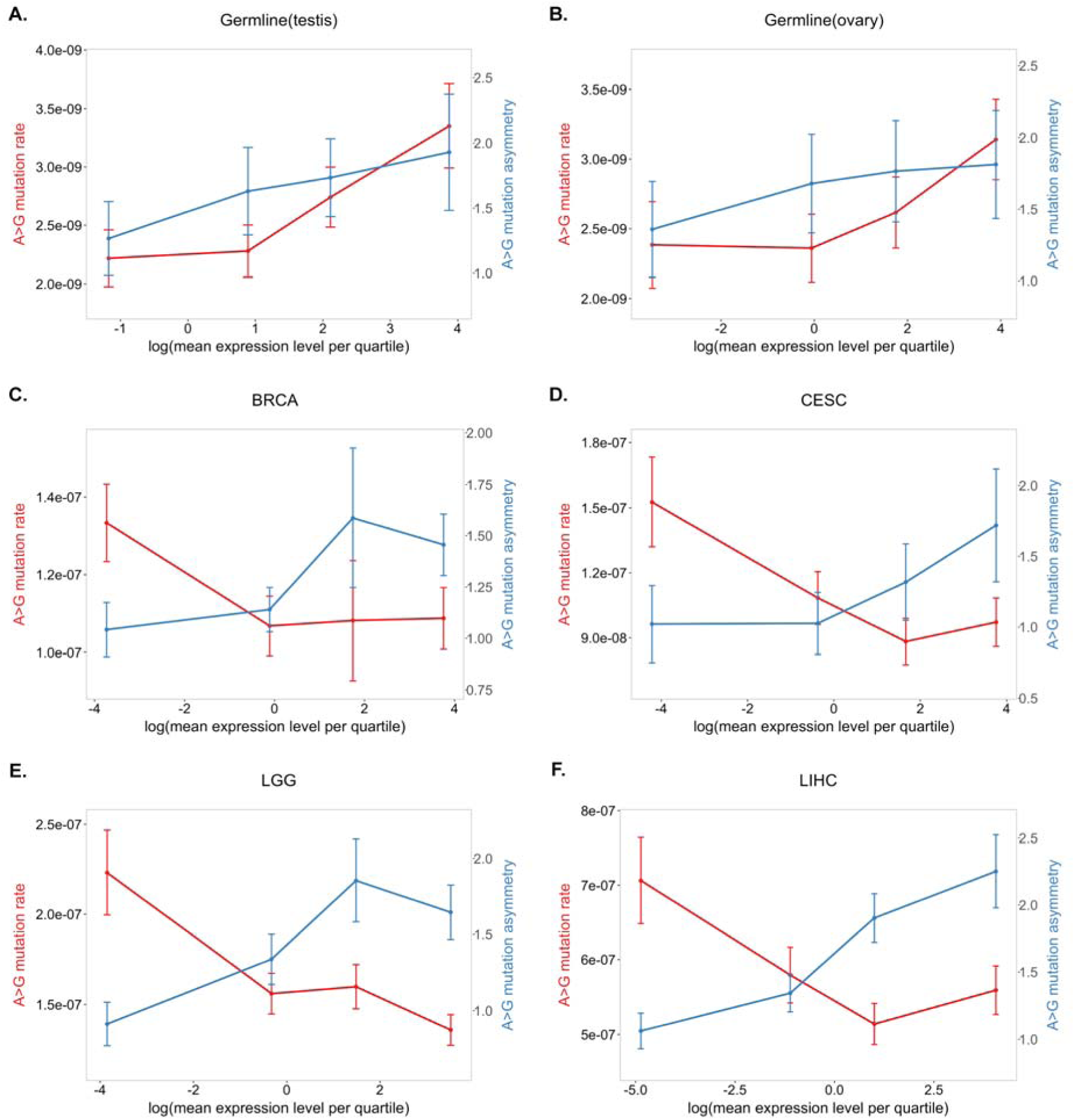
The degree of A>G strand asymmetry and the A>G mutation rate as a function of gene expression level quartiles. Shown are in panels A and B are results for the germline using testis expression levels and ovary expression levels, respectively; in panel C, for BRCA (breast invasive carcinoma); in panel D, for CESC (cervical squamous cell carcinoma and endocervical adenocarcinoma); in panel E, for LGG (brain lower grade glioma); and in panel F, for LIHC (liver hepatocellular carcinoma). The error bars for both the strand asymmetry and the mutation rate per quartile were estimated by bootstrapping (see Materials and Methods).

## Discussion

We compared the determinants of mutation in the soma and the germline, using the same unit of analysis (a coding region) and the same statistical model, and applied it to similar exome data for germline de novo mutations and four types of tumors, in which mutations largely predate tumorigenesis. We recapitulated previous findings of the effects of GC content and of a histone mark indicative of repression on germline and somatic mutations, as well as those of expression levels and replicating time on somatic mutations (Schuster-Böckler and Lehner 2012; Lawrence et al. 2013). Strikingly, we also found clear differences in the determinants of mutation rates between germline and soma, consistent with earlier hints based on divergence data (Hodgkinson and Eyre-Walker 2011). Notably, our results confirmed that somatic mutation rates decrease with expression levels and reveal that, in sharp contrast, de novo germline mutation rates increase with expression. This contrast suggests that transcription is mutagenic in germline but not in soma, and that the DNA damage or repair processes differ between them.

One limitation of our comparison—and of previous studies of germline and somatic mutation—is the need to rely on proxies for determinants of interest, such as replication timing data from cancer cell lines instead of normal cells. A second limitation is that we considered only two types of mutations (CpG Ti and other). Other work indicates that while these two types capture most of the variation in mutation rates, the larger context (adjacent base pairs, but also 7mers) also impacts mutation rates (Hwang and Green 2004; Hodgkinson and Eyre-Walker 2011; Aggarwala and Voight 2016). These different mutation subtypes are likely affected somewhat differently by the determinants considered here (Carlson et al. 2017). Despite these limitations, our work provides a framework to contrast possible determinants of mutation rates in soma and germline while controlling for some confounding effects, and results will only improve as data sets increase and the measurements of salient genomic and cellular features become more accurate. What is already clear is that the divergent effect of expression on mutation rates in germline and soma is not attributable to well-known covariates (included in our model). Moreover, the differences that cannot readily be explained by the noise introduced by imperfect proxies or limited data.

Notably, our results indicate that the tradeoff between damage and repair associated with transcription must differ between germline and soma. Transcription plausibly increases the rate of damage by opening up the DNA helix, rendering the single strands more susceptible to mutagens (Polak and Arndt 2008; Jinks-Robertson and Bhagwat 2014). One possibility is that, in the germline, the rate of transcription-associated mutagenesis (TAM) swamps TCR, leading to higher mutation rates with increased transcription, whereas in the soma, TCR is relatively more efficient and the balance of TAM and TCR leads to decreased mutagenesis with increased expression. Another possibility, which is not mutually exclusive, is the presence of additional repair mechanisms in somatic tissues. In support of this possibility, global genome repair (GGR) is attenuated in differentiated cells, yet mutations on the NTS appear to nonetheless be repaired efficiently (Nouspikel and Hanawalt 2000; Marteijn et al. 2014). This evidence led to the hypothesis of transcription-domain-associated repair (DAR), which might repair damage on both strands in addition to TCR (reviewed in Nouspikel 2007). From an evolutionary standpoint, the increased efficiency of TCR relative to TAM in soma versus germline may be explained by selection pressure on the repair of somatic tissues to prevent aging and cancer (Lynch 2010).

Mounting evidence suggests that per cell division mutation rates differ across tissues (Greenman et al. 2007; Lynch 2010; Alexandrov et al. 2013) and in particular that they may be higher in early embryonic development than at other stages of development (Ségurel, Wyman, and Przeworski 2014; Rahbari et al. 2016; Harland et al. 2016; Lindsay et al. 2016). This study further suggests that at least part of the explanation may lie in the balance between damage and repair, with TCR operating at different efficiencies relative to TAM or jointly with other repair pathways, thereby maintaining low mutation rates in soma. As mutation data from more tissues become available, it will be both feasible and enlightening to examine tissue-specific differences in repair.

## Acknowledgments

We thank Ziyue Gao and Priya Moorjani for comments on the manuscripts and helpful discussions.

## References

Aggarwala, Varun, and Benjamin F. Voight. 2016. “An Expanded Sequence Context Model Broadly Explains Variability in Polymorphism Levels across the Human Genome.” Nature Genetics 48 (4): 349–55. doi:10.1038/ng.3511.

Alexandrov, Ludmil B., Philip H. Jones, David C. Wedge, Julian E. Sale, Peter J. Campbell, Serena Nik-Zainal, and Michael R. Stratton. 2015. “Clock-like Mutational Processes in Human Somatic Cells.” Nature Genetics 47 (12): 1402–7. doi:10.1038/ng.3441.

Alexandrov, Ludmil, Nik-Zainal Serena, David Wedge, Samuel Aparicio, Sam Behjati, Andrew Biankin, Graham Bignell, et al. 2013. “Signatures of Mutational Processes in Human Cancer.” Nature 500 (7463): 415–21. doi:10.1038/nature12477.

Behjati, Sam, Meritxell Huch, Ruben van Boxtel, Wouter Karthaus, David C. Wedge, Asif U. Tamuri, Iñigo Martincorena, et al. 2014. “Genome Sequencing of Normal Cells Reveals Developmental Lineages and Mutational Processes.” Nature 513 (7518): 422–25. doi:10.1038/nature13448.

Besenbacher, Søren, Patrick Sulem, Agnar Helgason, Hannes Helgason, Helgi Kristjansson, Aslaug Jonasdottir, Adalbjorg Jonasdottir, et al. 2016. “Multi-Nucleotide de Novo Mutations in Humans.” PLOS Genetics 12 (11): e1006315. doi:10.1371/journal.pgen. 1006315.

Blokzijl, Francis, Joep de Ligt, Myrthe Jager, Valentina Sasselli, Sophie Roerink, Nobuo Sasaki, Meritxell Huch, et al. 2016. “Tissue-Specific Mutation Accumulation in Human Adult Stem Cells during Life.” Nature 538 (7624): 260–64. doi:10.1038/nature19768.

Bloom, L. B., M. R. Otto, R. Eritja, L. J. Reha-Krantz, M. F. Goodman, and J. M. Beechem. 1994. “Pre-Steady-State Kinetic Analysis of Sequence-Dependent Nucleotide Excision by the 3’-exonuclease Activity of Bacteriophage T4 DNA Polymerase.” Biochemistry 33 (24): 7576–86.

Brennan, Cameron W, Roel Verhaak, McKenna Aaron, Benito Campos, Houtan Noushmehr, Sofie R Salama, Siyuan Zheng, et al. 2013. “The Somatic Genomic Landscape of Glioblastoma.” 155 (2): 462–77. doi:10.1016/j.cell.2013.09.034.

Campbell, Catarina D, and Evan E Eichler. 2013. “Properties and Rates of Germline Mutations in Humans” 29 (10): 575–84. doi:10.1016/j.tig.2013.04.005.

Carlson, Jedidiah, Laura J. Scott, Adam E. Locke, Matthew Flickinger, Shawn Levy, The BRIDGES Consortium, Richard M. Myers, et al. 2017. “Extremely Rare Variants Reveal Patterns of Germline Mutation Rate Heterogeneity in Humans.” bioRxiv, February, 108290. doi:10.1101/108290.

De Rubeis, Silvia, Xin He, Arthur P. Goldberg, Christopher S. Poultney, Kaitlin Samocha, A. Ercument Cicek, Yan Kou, et al. 2014. “Synaptic, Transcriptional and Chromatin Genes Disrupted in Autism.” Nature 515 (7526): 209–15. doi:10.1038/nature13772.

Duret, Laurent, and Nicolas Galtier. 2009. “Biased Gene Conversion and the Evolution of Mammalian Genomic Landscapes.” Annual Review of Genomics and Human Genetics 10 (1): 285–311. doi:10.1146/annurev-genom-082908-150001.

Elango, Navin, Seong-Ho Kim, NISC Comparative Sequencing Program, Eric Vigoda, and Soojin V. Yi. 2008. “Mutations of Different Molecular Origins Exhibit Contrasting Patterns of Regional Substitution Rate Variation.” PLOS Computational Biology 4 (2): e1000015. doi:10.1371/journal.pcbi. 1000015.

Epi4K Consortium, and Epilepsy Phenome/Genome Project. 2013. “De Novo Mutations in Epileptic Encephalopathies.” Nature 501 (7466): 217–21. doi:10.1038/nature12439.

Francioli, Laurent C., Paz P. Polak, Amnon Koren, Androniki Menelaou, Sung Chun, Ivo Renkens, Genome of the Netherlands Consortium, et al. 2015. “Genome-Wide Patterns and Properties of de Novo Mutations in Humans.” Nature Genetics 47 (7): 822–26. doi:10.1038/ng.3292.

Fryxell, Karl J., and Won-Jong Moon. 2005. “CpG Mutation Rates in the Human Genome Are Highly Dependent on Local GC Content.” Molecular Biology and Evolution 22 (3): 65–58. doi:10.1093/molbev/msi043.

Gao, Ziyue, Minyoung J. Wyman, Guy Sella, and Molly Przeworski. 2016. “Interpreting the Dependence of Mutation Rates on Age and Time.” PLOS Biology 14 (1): e1002355. doi:10.1371/journal.pbio.1002355.

Goldmann, Jakob M, Wendy SW Wong, Michele Pinelli, Terry Farrah, Dale Bodian, Anna B Stittrich, Gustavo Glusman, et al. 2016. “Parent-of-Origin-Specific Signatures of de Novo Mutations” 48 (8): 935–39. doi:10.1038/ng.3597.

Green, Phil, Brent Ewing, Webb Miller, Pamela J. Thomas, NISC Comparative Sequencing Program, and Eric D. Green. 2003. “Transcription-Associated Mutational Asymmetry in Mammalian Evolution.” Nature Genetics 33 (4): 514–17. doi:10.1038/ng1103.

Greenman, Christopher, Philip Stephens, Raffaella Smith, Gillian L Dalgliesh, Christopher Hunter, Graham Bignell, Helen Davies, et al. 2007. “Patterns of Somatic Mutation in Human Cancer Genomes.” Cah Rev The 446 (7132): 153–58. doi:10.1038/nature05610.

Hamdan, Fadi F., Myriam Srour, Jose-Mario Capo-Chichi, Hussein Daoud, Christina Nassif, Lysanne Patry, Christine Massicotte, et al. 2014. “De Novo Mutations in Moderate or Severe Intellectual Disability.” PLOS Genetics 10 (10): e1004772. doi:10.1371/journal.pgen.1004772.

Hanawalt, Philip C, and Graciela Spivak. 2008. “Transcription-Coupled DNA Repair: Two Decades of Progress and Surprises” 9 (12): 958–70. doi:10.1038/nrm2549.

Harland, Chad, Carole Charlier, Latifa Karim, Nadine Cambisano, Manon Deckers, Erik Mullaart, Wouter Coppieters, and Michel Georges. 2016. “Frequency of Mosaicism Points towards Mutation-Prone Early Cleavage Cell Divisions.” bioRxiv October, 79863. doi:10.1101/079863.

Hodgkinson, Alan, Ying Chen, and Adam Eyre-Walker. 2012. “The Large□scale Distribution of Somatic Mutations in Cancer Genomes” 33 (1): 136–43. doi:10.1002/humu.21616.

Hodgkinson, Alan, and Adam Eyre-Walker. 2011. “Variation in the Mutation Rate across Mammalian Genomes.” Nat Rev Genetics 12 (11): 756–66. doi:10.1038/nrg3098.

Homsy, Jason, Samir Zaidi, Yufeng Shen, James S. Ware, Kaitlin E., Samocha, Konrad J. Karczewski, Steven R. DePalma, et al. 2015. “De Novo Mutations in Congenital Heart Disease with Neurodevelopmental and Other Congenital Anomalies.” Science 350 (6265): 1262–66. doi:10.1126/science.aac9396.

Hwang, Dick G., and Phil Green. 2004. “Bayesian Markov Chain Monte Carlo Sequence Analysis Reveals Varying Neutral Substitution Patterns in Mammalian Evolution.” Proceedings of the National Academy of Sciences of the United States of America 101 (39): 13994–1. doi:10.1073/pnas. 0404142101.

Iossifov, Ivan, Brian J. O’Roak, Stephan J. Sanders, Michael Ronemus, Niklas Krumm, Dan Levy, Holly A. Stessman, et al. 2014. “The Contribution of deovo Coding Mutations to Autism Spectrum Disorder.” Nature 515 (7526): 216–21. doi:10.1038/nature13908.

Jinks-Robertson, Sue, and Ashok S. Bhagwat. 2014. “Transcription-Associated Mutagenesis.” Annual Review of Genetics 48 (1): 341–59. doi:10.1146/annurev-genet-120213-092015.

Kong, Augustine, Michael L. Frigge, Gisli Masson, Soren Besenbacher, Patrick Sulem, Gisli Magnusson, Sigurjon A. Gudjonsson, et al. 2012. “Rate of de Novo Mutations and the Importance of Father/’s Age to Disease Risk.” Nature 488 (7412): 471–75. doi:10.1038/nature11396.

Koren, Amnon, Paz Polak, James Nemesh, Jacob J. Michaelson, Jonathan Sebat, Shamil R. Sunyaev, and Steven A. McCarroll. 2012. “Differential Relationship of DNA Replication Timing to Different Forms of Human Mutation and Variation.” The American Journal of Human Genetics 91 (6): 1033–40. doi:10.1016/j.ajhg.2012.10.018.

Larsson, Thomas P., Christian G. Murray, Tobias Hill, Robert Fredriksson, and Helgi B. Schiöth. 2005. “Comparison of the Current RefSeq, Ensembl and EST Databases for Counting Genes and Gene Discovery.” FEBS Letters 579 (3): 690–98. doi:10.1016/j.febslet.2004.12.046.

Lawrence, Michael S, Petar Stojanov, Paz Polak, Gregory V Kryukov, Kristian Cibulskis, Andrey Sivachenko, Scott L Carter, et al. 2013. “Mutational Heterogeneity in Cancer and the Search for New Cancer-Associated Genes.” Nature 499 (7457): 214–18. doi:10.1038/nature12213.

Lee, William, Zhaoshi Jiang, Jinfeng Liu, Peter M. Haverty, Yinghui Guan, Jeremy Stinson, Peng Yue, et al. 2010. “The Mutation Spectrum Revealed by Paired Genome Sequences from a Lung Cancer Patient.” Nature 465 (7297): 473–77. doi:10.1038/nature09004.

Ligt, Joep de, Marjolein H. Willemsen, Bregje W.M. van Bon, Tjitske Kleefstra, Helger G. Yntema, Thessa Kroes, Anneke T. Vulto-van Silfhout, et al. 2012. “Diagnostic Exome Sequencing in Persons with Severe Intellectual Disability.” New England Journal of Medicine 367 (20): 1921–29. doi:10.1056/NEJMoa1206524.

Lindsay, Sarah J., Raheleh Rahbari, Joanna Kaplanis, Thomas Keane, and Matthew Hurles. 2016. “Striking Differences in Patterns of Germline Mutation between Mice and Humans.” bioRxiv, October, 82297. doi:10.1101/082297.

Lodato, Michael A, Mollie B Woodworth, Semin Lee, Gilad D Evrony, Bhaven K Mehta, Amir Karger, Soohyun Lee, et al. 2015. “Somatic Mutation in Single Human Neurons Tracks Developmental and Transcriptional History” 350 (6256): 94–98. doi:10.1126/science.aab1785.

Lynch, Michael. 2010. “Rate, Molecular Spectrum, and Consequences of Human Mutation.” Proc National Acad Sci 107 (3): 961–68. doi:10.1073/pnas.0912629107.

Marteijn, Jurgen A., Hannes Lans, Wim Vermeulen, and Jan H. J. Hoeijmakers. 2014. “Understanding Nucleotide Excision Repair and Its Roles in Cancer and Ageing.” Nature Reviews Molecular Cell Biology 15 (7): 465–81. doi:10.1038/nrm3822.

Martincorena, Iñigo, Amit Roshan, Moritz Gerstung, Peter Ellis, Peter Loo, McLaren Stuart, David C Wedge, et al. 2015. “High Burden and Pervasive Positive Selection of Somatic Mutations in Normal Human Skin” 348 (6237): 880–86. doi:10.1126/science.aaa6806.

McVicker, Graham, David Gordon, Colleen Davis, and Phil Green. 2009. “Widespread Genomic Signatures of Natural Selection in Hominid Evolution.” PLOS Genetics 5 (5): e1000471. doi:10.1371/journal.pgen.1000471.

McVicker, Graham, and Phil Green. 2010. “Genomic Signatures of Germline Gene Expression.” Genome Research 20 (11): 1503–11. doi:10.1101/gr.106666.110.

Michaelson, Jacob J, Yujian Shi, Madhusudan Gujral, Hancheng Zheng, Dheeraj Malhotra, Xin Jin, Minghan Jian, et al. 2012. “Whole-Genome Sequencing in Autism Identifies Hot Spots for De Novo Germline Mutation” 151 (7): 1431–42. doi:10.1016/j.cell.2012.11.019.

Moorjani, Priya, Ziyue Gao, and Molly Przeworski. 2016. “Human Germline Mutation and the Erratic Evolutionary Clock.” PLOS Biology 14 (10): e2000744. doi:10.1371/journal.pbio.2000744.

Muller, H. J. 1927. “ARTIFICIAL TRANSMUTATION OF THE GENE.” Science 66 (1699): 84–87. doi:10.1126/science.66.1699.84.

Nouspikel, Thierry. 2007. “DNA Repair in Differentiated Cells: Some New Answers to Old Questions.” Neuroscience, Genome Dynamics and DNA Repair in the CNS, 145 (4): 1213–21. doi:10.1016/j.neuroscience.2006.07.006.

Nouspikel, Thierry. 2009. “DNA Repair in Mammalian Cells.” Cellular and Molecular Life Sciences 66 (6): 994–1009. doi:10.1007/s00018-009-8737-y.

Nouspikel, Thierry, and Philip C. Hanawalt. 2000. “Terminally Differentiated Human Neurons Repair Transcribed Genes but Display Attenuated Global DNA Repair and Modulation of Repair Gene Expression.” Molecular and Cellular Biology 20 (5): 1562–70.

Park, Chungoo, Wenfeng Qian, and Jianzhi Zhang. 2012. “Genomic Evidence for Elevated Mutation Rates in Highly Expressed Genes.” EMBO Reports 13 (12): 1123–29. doi:10.1038/embor.2012.165.

Petruska, J., and M. F. Goodman. 1985. “Influence of Neighboring Bases on DNA Polymerase Insertion and Proofreading Fidelity.” Journal of Biological Chemistry 260 (12): 7533–39.

Pleasance, Erin D., R. Keira Cheetham, Philip J. Stephens, David J. McBride, Sean J. Humphray, Chris D. Greenman, Ignacio Varela, et al. 2010. “A Comprehensive Catalogue of Somatic Mutations from a Human Cancer Genome.” Nature 463 (7278): 191–96. doi:10.1038/nature08658.

Pleasance, Erin D., Philip J. Stephens, Sarah O’Meara, David J. McBride, Alison Meynert, David Jones, Meng-Lay Lin, et al. 2010. “A Small-Cell Lung Cancer Genome with Complex Signatures of Tobacco Exposure.” Nature 463 (7278): 184–90. doi:10.1038/nature08629.

Polak, Paz, and Peter F. Arndt. 2008. “Transcription Induces Strand-Specific Mutations at the 5' End of Human Genes.” Genome Research 18 (8): 1216–23. doi:10.1101/gr.076570.108.

Polak, Paz, Rosa Karlić, Amnon Koren, Robert Thurman, Richard Sandstrom, Michael Lawrence, Alex Reynolds, et al. 2015. “Cell-of-Origin Chromatin Organization Shapes the Mutational Landscape of Cancer.” Nature 518 (7539): 360–64. doi:10.1038/nature14221.

Rahbari, Raheleh, Arthur Wuster, Sarah Lindsay, Robert Hardwick, Ludmil Alexandrov, Saeed Turki, Anna Dominiczak, et al. 2016. “Timing, Rates and Spectra of Human Germline Mutation.” Nat Genet 48 (2): 126–33. doi:10.1038/ng.3469.

Rauch, Anita, Dagmar Wieczorek, Elisabeth Graf, Thomas Wieland, Sabine Endele, Thomas Schwarzmayr, Beate Albrecht, et al. 2012. “Range of Genetic Mutations Associated with Severe Non-Syndromic Sporadic Intellectual Disability: An Exome Sequencing Study.” The Lancet 380 (9854): 1674–82. doi:10.1016/S0140-6736(12)61480-9.

Rubin, Alan F., and Phil Green. 2009. “Mutation Patterns in Cancer Genomes.” Proceedings of the National Academy of Sciences 106 (51): 21766–70. doi:10.1073/pnas.0912499106.

Samocha, Kaitlin E, Elise B Robinson, Stephan J Sanders, Christine Stevens, Aniko Sabo, Lauren M McGrath, Jack A Kosmicki, et al. 2014. “A Framework for the Interpretation of de Novo Mutation in Human Disease.” Nature Genetics 46 (9): 944–50. doi:10.1038/ng.3050.

Schuster-Böckler, Benjamin, and Ben Lehner. 2012. “Chromatin Organization Is a Major Influence on Regional Mutation Rates in Human Cancer Cells” 488 (7412): 504–7. doi:10.1038/nature11273.

Ségurel, Laure, Minyoung J. Wyman, and Molly Przeworski. 2014. “Determinants of Mutation Rate Variation in the Human Germline.” Annual Review of Genomics and Human Genetics 15: 47–70. doi:10.1146/annurev-genom-031714-125740.

Shendure, Jay, and Joshua M. Akey. 2015. “The Origins, Determinants, and Consequences of Human Mutations.” Science 349 (6255): 1478–83. doi:10.1126/science.aaa9119.

Stamatoyannopoulos, John A., Ivan Adzhubei, Robert E. Thurman, Gregory V. Kryukov, Sergei M. Mirkin, and Shamil R. Sunyaev. 2009. “Human Mutation Rate Associated with DNA Replication Timing.” Nature Genetics 41 (4): 393–95. doi:10.1038/ng.363.

Stratton, Michael R., 2011. “Exploring the Genomes of Cancer Cells: Progress and Promise.” Science 331 (6024): 1553–58. doi:10.1126/science.1204040.

Stratton, Michael R., Peter J. Campbell, and P. Andrew Futreal. 2009. “The Cancer Genome.” Nature 458 (7239): 719–24. doi:10.1038/nature07943.

Supek, Fran, and Ben Lehner. 2015. “Differential DNA Mismatch Repair Underlies Mutation Rate Variation across the Human Genome” 521 (7550): 81–84. doi:10.1038/nature14173.

Takai, Daiya, and Peter A. Jones. 2002. “Comprehensive Analysis of CpG Islands in Human Chromosomes 21 and 22.” Proceedings of the National Academy of Sciences 99 (6): 3740–45. doi:10.1073/pnas.052410099.

The Deciphering Developmental Disorders Study. 2015. “Large-Scale Discovery of Novel Genetic Causes of Developmental Disorders.” Nature 519 (7542): 223–28. doi:10.1038/nature14135.

Webster, Matthew T., Nick G. C. Smith, Martin J. Lercher, and Hans Ellegren. 2004. “Gene Expression, Synteny, and Local Similarity in Human Noncoding Mutation Rates.” Molecular Biology and Evolution 21 (10): 1820–30. doi:10.1093/molbev/msh181.

Zhao, Shanrong, and Baohong Zhang. 2015. “A Comprehensive Evaluation of Ensembl, RefSeq, and UCSC Annotations in the Context of RNA-Seq Read Mapping and Gene Quantification.” BMC Genomics 16 (1): 97. doi:10.1186/s12864-015-1308-8.

